# Local induction of *IAA5* and *IAA29* promotes DNA damage-triggered stem cell death in *Arabidopsis* roots

**DOI:** 10.1101/2022.09.02.506394

**Authors:** Naoki Takahashi, Nobuo Ogita, Toshiya Koike, Kohei Nishimura, Soichi Inagaki, Masaaki Umeda

**Author notes:** To whom correspondence should be addressed: Masaaki Umeda, Graduate School of Science and Technology, Nara Institute of Science and Technology, Takayama 8916-5, Ikoma, Nara 630-0192, Japan. Department of Biological Sciences, Graduate School of Science, The University of Tokyo, Hongo, Bunkyo-ku, Tokyo 113-0033, Japan (Soichi Inagaki). The author responsible for distribution of materials integral to the findings presented in this article in accordance with the policy described in the Instructions for Authors (https://academic.oup.com/plphys/pages/General-Instructions) is: Masaaki Umeda. **Author contributions** N.T., and M.U. designed the research; N.T., O.N., T.K., K.N., and S.I. performed the experiments and analyzed data; N.T. and M.U. wrote the article; and M.U. supervised the research.

## Abstract

Plants generate organs continuously during post-embryonic development, thus how to preserve stem cells in changing environments is crucial for their survival. Genotoxic stress threatens genome stability in all somatic cells, whereas, in the meristem, only stem cells undergo cell death in response to DNA damage, followed by stem cell replenishment. It was previously shown that inhibition of downward auxin flow in roots participates in DNA damage-induced stem cell death; however, how cell death is confined to stem cells in tissues with reduced auxin content remains elusive. Here we show that, in response to DNA double-strand breaks, the Aux/IAA family members *IAA5* and *IAA29*, which encode the negative regulators of auxin signaling, were locally induced in vascular stem cells and their daughters in *Arabidopsis* roots. This is an active response to DNA damage in which the transcription factor SUPPRESSOR OF GAMMA RESPONSE 1 directly induces their expression. In the *iaa5 iaa29* double mutant, DNA damage-induced stem cell death was significantly suppressed, while it was fully restored by the expression of a stable form of IAA5 in vascular stem cells. Our genetic data revealed that both the suppression of auxin signaling and reduced auxin content are prerequisite for cell death induction, suggesting the central role in maintaining genome integrity in stem cells.

**One-sentence summary:** The induction of *IAA5* and *IAA29* promotes stem cell death in *Arabidopsis* roots under DNA stress.

## Introduction

Maintenance of genome integrity is crucial for survival of any organism. Especially, in plants, stem cells need to preserve genome stability throughout their lives to achieve continuous organ growth and development. DNA is damaged by internal factors, such as DNA replication errors and reactive oxygen species, but also by external factors, e.g., ultraviolet and gamma irradiation and oxidative stress (Ciccia and Elledge, 2010). Previous studies demonstrated that high boron or aluminum stress, as well as pathogen infection also cause DNA damage in plants (Rounds and Larsen, 2008; Sakamoto et al., 2011; Song and Bent, 2014). In eukaryotes, DNA damage is sensed by the two kinases ATAXIA-TELANGIECTASIA MUTATED (ATM) and ATM/RAD3-RELATED (ATR) (Maréchal and Zou, 2013): ATM is activated by DNA double-strand breaks (DSBs), whereas ATR senses stalled replication forks and single-strand breaks (SSBs). However, in plants, downstream regulators that are orthologous to mammalian counterparts are all missing, and the plant-specific NAC-type transcription factor SUPPRESSOR OF GAMMA RESPONSE 1 (SOG1) plays a crucial role in transmitting DNA damage signals (Yoshiyama et al., 2009). SOG1 is phosphorylated and activated by ATM and ATR (Yoshiyama et al., 2013; Sjogren et al., 2015), and a recent study showed that it is stabilized through monoubiquitination by the E3 ligase DDRM1 (Wang et al., 2022). SOG1 binds to the consensus sequence CTT[N]_7_AAG and induces the expression of target genes involved in DNA repair, cell cycle arrest, and plant immunity (Bourbousse et al., 2018; Ogita et al., 2018).

In the *Arabidopsis* roots tip, stem cells surround the quiescent center (QC), which has very low mitotic activity and is involved in stem cell maintenance (van den Berg et al., 1997; Ortega-Martínez et al., 2007). The descendants of stem cells divide several times in the meristematic zone (MZ) and eventually stop cell division to start endoreplication, a repeated cycle of DNA replication without mitosis or cytokinesis. Endoreplicating cells initiate active cell elongation in the transition zone (TZ), followed by more rapid cell elongation in the elongation/differentiation zone (EDZ), which is triggered by actin reorganization (Takatsuka et al., 2018). We previously reported that DSBs arrest the cell cycle at G2 phase in the MZ, and promote an early onset of endoreplication at the boundary between the MZ and the TZ (Adachi et al., 2011; Chen et al., 2017). Another prominent feature of the plant DNA damage response is stem cell death. While cell death is generally observed in animal cells exposed to severe DNA damage, only stem cells are going to die in the shoot and root meristems (Fulcher and Sablowski, 2009). These observations indicate that different types of events occur in distinct zones of roots, thereby balancing genome maintenance with continuous root growth.

We previously found that SOG1 directly induces the NAC-type transcription factors ANAC044 and ANAC085, and stabilizes R1R2R3-Myb transcription factors MYB3R3 and MYB3R5 that repress a set of G2/M-specific genes, resulting in G2 arrest (Chen et al., 2017; Takahashi et al., 2019). The protein stability of the repressor-type R1R2R3-Myb transcription factors (Rep-MYBs) is also controlled by cyclin-dependent kinase (CDK) activity; in dividing cells, CDK phosphorylates and destabilizes Rep-MYBs, whereas DNA damage reduces CDK activities, thereby decreasing the phosphorylation level and causing Rep-MYB accumulation (Chen et al., 2017). However, as described above, ANAC044/085 play an essential role in Rep-MYB stabilization under DNA stress, thus it is likely that the SOG1-ANAC044/085 pathway inhibits protein degradation machineries and/or regulates Rep-MYB-interacting factors that affect Rep-MYB stability. Interestingly, in the root tip of the *anac044, anac085*, or *rep-myb* mutant, not only G2 arrest but also stem cell death is suppressed under DNA damage conditions (Chen et al., 2017; Takahashi et al., 2019), suggesting that G2 arrest is one of the requirements for stem cell death induction.

We have recently reported that DSBs upregulate the expression of cytokinin biosynthesis genes, and increase the cytokinin level in the *Arabidopsis* root tip (Takahashi et al., 2021). This hormonal response promotes two cytokinin-dependent events, an onset of endoreplication and a reduction of the auxin content. The cytokinin signaling activates the type-B response regulator ARR2, which then induces *CCS52A1*, encoding an activator of the anaphase-promoting complex/cyclosome (APC/C). The E3 ubiquitin ligase APC/C targets mitotic cyclins, thereby arresting the cell cycle and promoting the transition to endoreplication (Takahashi et al., 2013). On the other hand, the type-B response regulators ARR1 and ARR12 induce the Aux/IAA family member *SHY2*, which represses the expression of a few *PIN-FORMED* (*PIN*) genes encoding auxin efflux carriers (Dello Ioio et al., 2008). Consequently, the auxin level decreases in the root tip, reducing mitotic activity. Our data showed that exogenous auxin treatment dramatically suppressed DSB-induced G2 arrest and stem cell death (Takahashi et al., 2021), implying that cytokinin-dependent reduction of the auxin level is one of the major causes of the DNA damage response. However, it remains elusive how cell death is confined to stem cells although auxin reduction occurs in a wider area of the root meristem.

In this study, we identified two Aux/IAA family members *IAA5* and *IAA29*, whose expression is directly induced by SOG1. Both *IAA*s are scarcely expressed under normal growth conditions, while they are highly induced by DSBs in vascular stem cells and their daughters in the root tip. Our genetic analysis revealed that, in addition to the *SHY2*-dependent pathway, *IAA5*- and *IAA29*-mediated signaling contributes to a reduction in auxin signaling and causes stem cell death under DNA stress, highlighting the role of Aux/IAA family members in the maintenance of plant genome integrity.

## Results

### DSBs induce *IAA5* and *IAA29* through the SOG1-dependent pathway

We previously reported that DSBs upregulate cytokinin biosynthesis, resulting in decreased auxin level in the root tip (Takahashi et al., 2021). This is an indirect regulation of auxin signaling; therefore, we first investigated transcriptomic changes of auxin-related genes to know whether there is any direct mechanism controlling auxin signaling under DNA stress. We examined our previous microarray data, which were obtained from two-week-old *Arabidopsis* seedlings treated with the DSB inducer zeocin for 2 h, and quantified the mRNA levels of genes for auxin biosynthesis, perception, and signaling (Ogita et al., 2018). The dataset showed that, after zeocin treatment, no significant changes were observed in genes for auxin biosynthesis (*TAA1, TAR*s, and *YUCCA*s), auxin receptors (*TIR1/AFB*s), or auxin response factors (*ARF*s) (Supplemental Figure S1, A–C). However, among the 31 genes encoding the Aux/IAA family members, *IAA5* and *IAA29* exhibited a dramatic increase in transcript levels by zeocin treatment (Supplemental Figure S1D). The other datasets reported by Bourbousse et al. (2018) and Takahashi et al. (2019) also indicated that their transcript levels were elevated in response to DSBs caused by gamma irradiation or bleomycin treatment (Supplemental Figure S2).

To confirm the above-mentioned transcriptomic data, we conducted quantitative reverse transcription PCR (qRT-PCR). Five-day-old wild-type (WT) seedlings were transferred onto a Murashige and Skoog (MS) medium containing 0.6 μg/ml bleomycin, and grown for 0, 6, 12, and 24 h. Total RNA was extracted from the roots and subjected to qRT-PCR analysis. The result showed that the *IAA5* and *IAA29* transcripts already increased at 6 h and were gradually elevated afterward, indicating that they rapidly respond to DSBs (Figure 1A). Importantly, such an increase was not observed in the *sog1-101* knockout mutant (Figure 1A), implying a programmed response to DNA damage. Note that DSB-induced stem cell death, which was identified by staining with propidium iodide (PI), was visible 12–18 h after bleomycin treatment (Figure 1B), suggesting that *IAA5* and *IAA29* induction precedes stem cell death.

**Figure 1.**
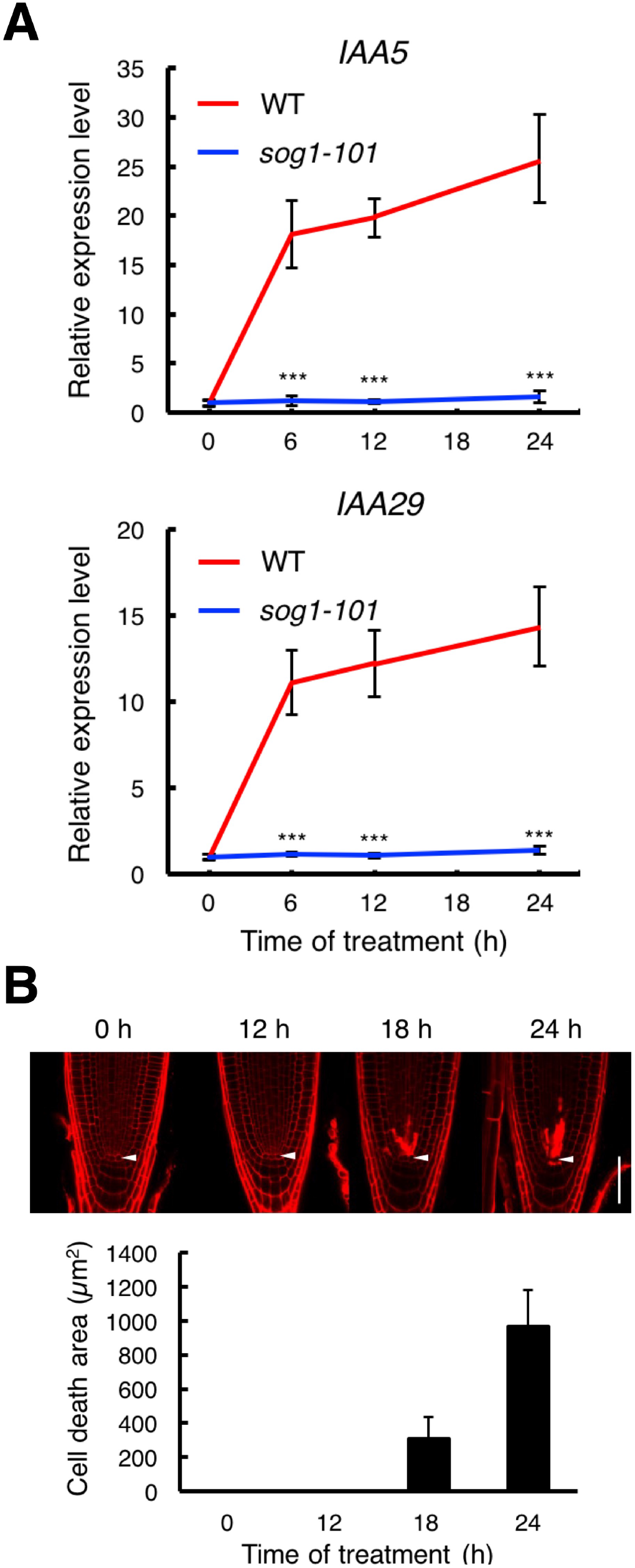
Bleomycin treatment induces *IAA5* and *IAA29* before the appearance of stem cell death. (A) Transcript levels of *IAA5* and *IAA29* in the presence of bleomycin. Five-day-old WT and *sog1-101* seedlings were transferred onto MS plates containing 0.6 μg/ml bleomycin, and grown for 0, 6, 12, and 24 h. Total RNA was extracted from roots and subjected to qRT-PCR. Transcript levels of *IAA5* and *IAA29* were normalized to that of *ACTIN2*, and are indicated as relative values, with that for 0 h set to 1. Data are presented as mean ± SD calculated from three biological replicates. For the data of *sog1-101*, significant differences from WT were determined by Student’s *t*-test: ***, *P* < 0.001. (B) Time-course observation of bleomycin-induced stem cell death. Five-day-old WT seedlings were transferred onto an MS medium containing 0.6 μg/ml bleomycin, and grown for 0, 12, 18, and 24 h. Root tips were stained with PI, and the area of PI-stained dead cells was measured using Fiji imaging software. Data are presented as mean ± SD (n > 20). Arrowheads indicate the QC. Bar = 100 μm.

### *IAA5* and *IAA29* are specifically induced in vascular stem cells and their daughters in response to DSBs

Next, we examined the spatial expression patterns of *IAA5* and *IAA29* in the root tip. We generated the transgenic plants harboring the *pIAA5:GFP* or *pIAA29:GFP* reporter genes, in which *GFP* is expressed under the 2-kb promoter of each *IAA*. Five-day-old plants were transferred onto a medium containing 0.6 μg/ml bleomycin, and GFP fluorescence was observed after 24 h. As shown in Figure 2, almost no fluorescence was detected in the root tip of both reporter lines under normal growth conditions. However, after 24-h-bleomycin treatment, a dramatic induction was observed in vascular stem cells and their daughters, which preferentially die in response to DSBs. Consistent with the qRT-PCR data, *IAA5* and *IAA29* expression was observed before stem cell death occurred, that is, after 12-h-bleomycin treatment (Supplemental Figure S3), and both reporter genes showed no GFP signal in *sog1-101* irrespective of the presence of bleomycin (Figure 2).

**Figure 2.**
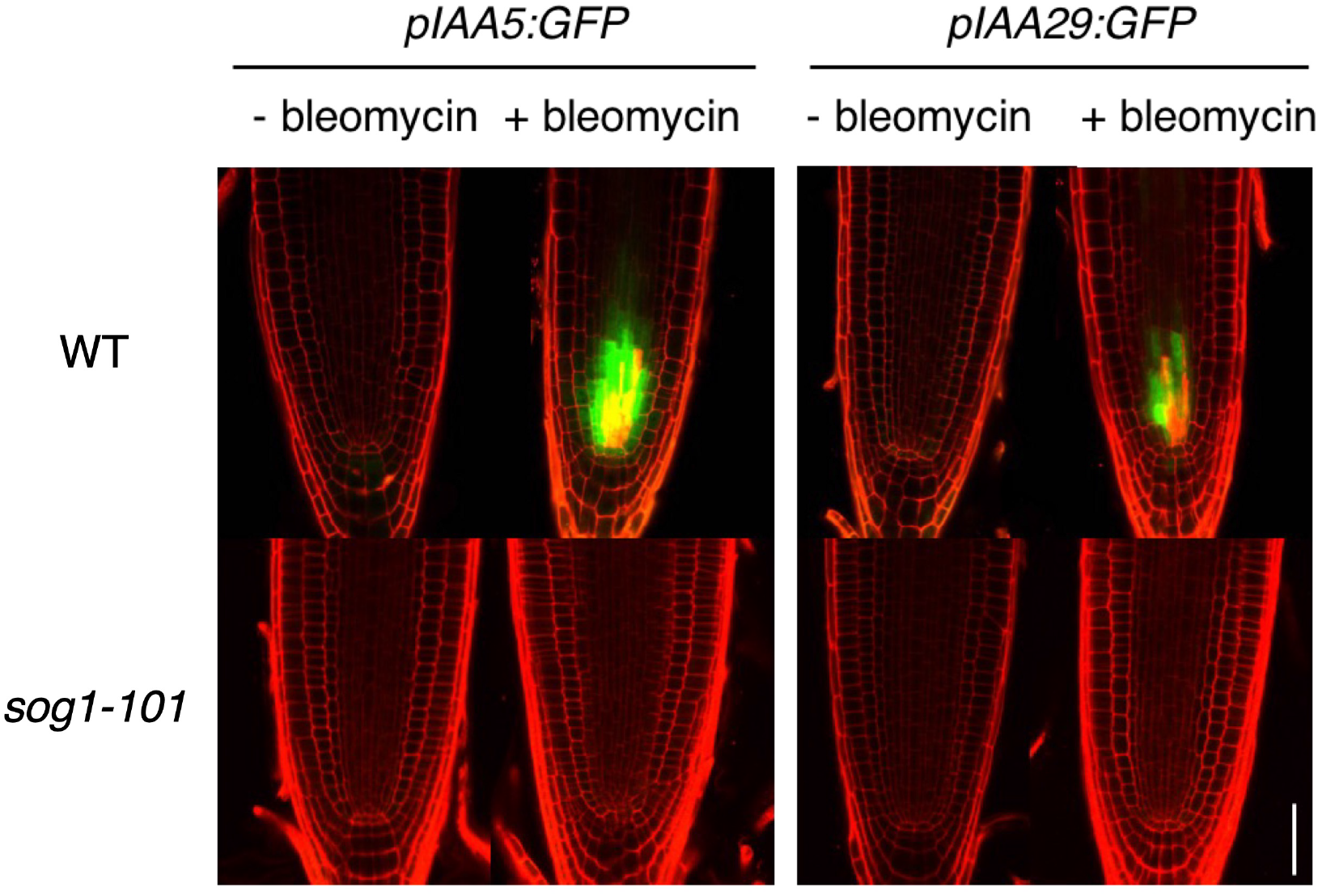
Local induction of *IAA5* and *IAA29* in bleomycin-treated root tips. Five-day-old WT and *sog1-101* seedlings harboring *pIAA5:GFP* or *pIAA29:GFP* were transferred onto MS plates with or without 0.6 μg/ml bleomycin, and grown for 24 h. GFP fluorescence was observed after counterstaining with PI. Bar = 100 μm.

The above-mentioned result suggests that DSBs, but not resultant stem cell death, induce *IAA5* and *IAA29*. However, a possibility still exists that an early invisible process leading to cell death upregulates *IAA5* and *IAA29*. To test this possibility, we conducted laser ablation of the stem cell region in *pIAA5:GFP* and *pIAA29:GFP*, and monitored GFP fluorescence after 0, 6, 12, 24 h. As a result, no fluorescence was observed in the root tip at any time points (Supplemental Figure S4), suggesting that *IAA5* and *IAA29* induction is not a consequence of stem cell death but requires the DSB signal.

### SOG1 binds to the promoter regions of *IAA5* and *IAA29* in response to DSBs

As described above, SOG1 is essential for *IAA5* and *IAA29* induction in response to DSBs (Figures 1A and 2); therefore, we examined the possibility that *IAA5* and *IAA29* are direct targets of SOG1. Previous chromatin immunoprecipitation (ChIP) sequence data show significant peaks of SOG1-bound gnomic DNA fragments in the promoter regions of *IAA5* and *IAA29* (Bourbousse et al., 2018; Ogita et al., 2018). In addition, the promoters harbor <u>CTT</u>GTCTCAA<u>ACG</u> and <u>CGT</u>GTTGGTT<u>AAG</u> sequences, respectively, which are closely related to the SOG-binding consensus sequence CTT(N)_7_AAG (Ogita et al., 2018). These findings prompted us to perform ChIP-qPCR analysis of the *IAA5* and *IAA29* genes. ChIP was conducted with the transgenic plant harboring *pSOG1:SOG1-Myc* (Yoshiyama et al., 2013), and genomic fragments bound to the anti-Myc antibody were subjected to qPCR using primers that amplify the regions of the promoter, gene body and 3′-UTR (Figure 3A). When *pSOG1:SOG1-Myc* seedlings were not treated with bleomycin, no SOG1 binding was detected in any region tested in this study (Figure 3B). However, genomic DNA prepared from bleomycin-treated plants displayed a high level of binding to the promoter, and a lower but significant level to the gene body and 3′-UTR of *IAA5* and *IAA29* (Figure 3B). Note that SOG1 requires DNA damage-triggered phosphorylation to efficiently bind to target sites (Ogita et al., 2018), supporting the above result of bleomycin-dependent SOG1 binding. These data indicate that SOG1 directly induces *IAA5* and *IAA29* in response to DNA damage.

**Figure 3.**
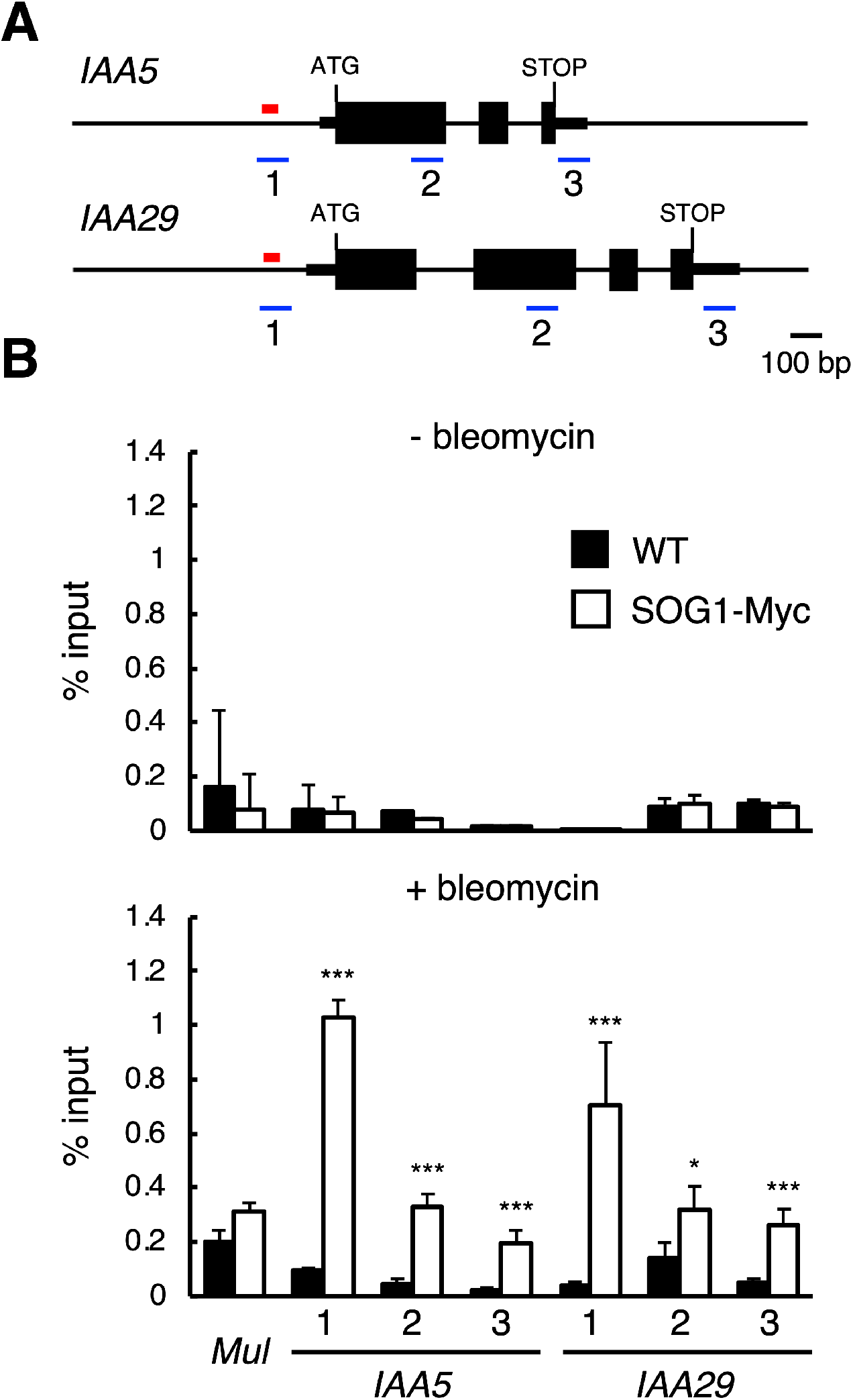
SOG1 binds to the genomic regions of *IAA5* and *IAA29*. (A) Schematic representation of the genomic regions of *IAA5* and *IAA29*. Red lines and black boxes indicate the SOG1-binding motifs and exons, respectively. Thin black boxes mark the untranslated regions (UTRs). The blue lines represent the regions amplified in the ChIP-qPCR assay. (B) ChIP-qPCR assay. Two-week-old WT and *pSOG1:SOG1-Myc* (SOG1-Myc) seedlings grown in a liquid MS medium were treated with or without 0.6 μg/ml bleomycin for 12 h. Chromatin bound to SOG1-Myc extracted from whole seedlings was collected by immunoprecipitation with anti-Myc antibodies, and qPCR was conducted to amplify the regions in the promoter (1), gene body (2), and 3′-UTR (3) of *IAA5* and *IAA29. Mutator-like transposon* (*Mul*) was used as a negative control. The recovery rate of each DNA fragment was determined against the input DNA. Data are presented as mean ± SD (n = 3). For the data of SOG1-Myc, significant differences from WT were determined by Student’s *t*-test: *, *p* < 0.05; ***, *p* < 0.001.

### *IAA5* and *IAA29* are redundantly involved in DSB-induced stem cell death

We recently reported that a reduction in auxin signaling in the root tip is involved in DNA damage-induced stem cell death (Takahashi et al., 2021). Considering that IAA5 and IAA29 function in reducing auxin signaling (Tiwari et al., 2001), their induction is likely associated with stem cell death. To test this hypothesis, we observed the response of the *iaa5-1* and *iaa29-1* knockout mutants to DSBs (Overvoorde et al., 2005; Jiang et al., 2014). Five-day-old seedlings were transferred onto an MS plate containing 0.6 μg/ml bleomycin, and the roots were stained with PI after 24 h. As shown in Figure 4, cell death area was almost the same between WT and each single mutant, whereas it was dramatically reduced in the *iaa5-1 iaa29-1* double mutant. This indicates that *IAA5* and *IAA29* are involved in DSB-induced stem cell death, and that they play redundant roles in reducing auxin signaling under DNA stress. Notably, in bleomycin-treated double mutants, cell death was suppressed but not completely (Figure 4), suggesting that other mechanism(s), such as cytokinin-dependent inhibition of *PIN* expression (Takahashi et al., 2021), also participate in decreasing auxin signaling and/or inducing stem cell death.

**Figure 4.**
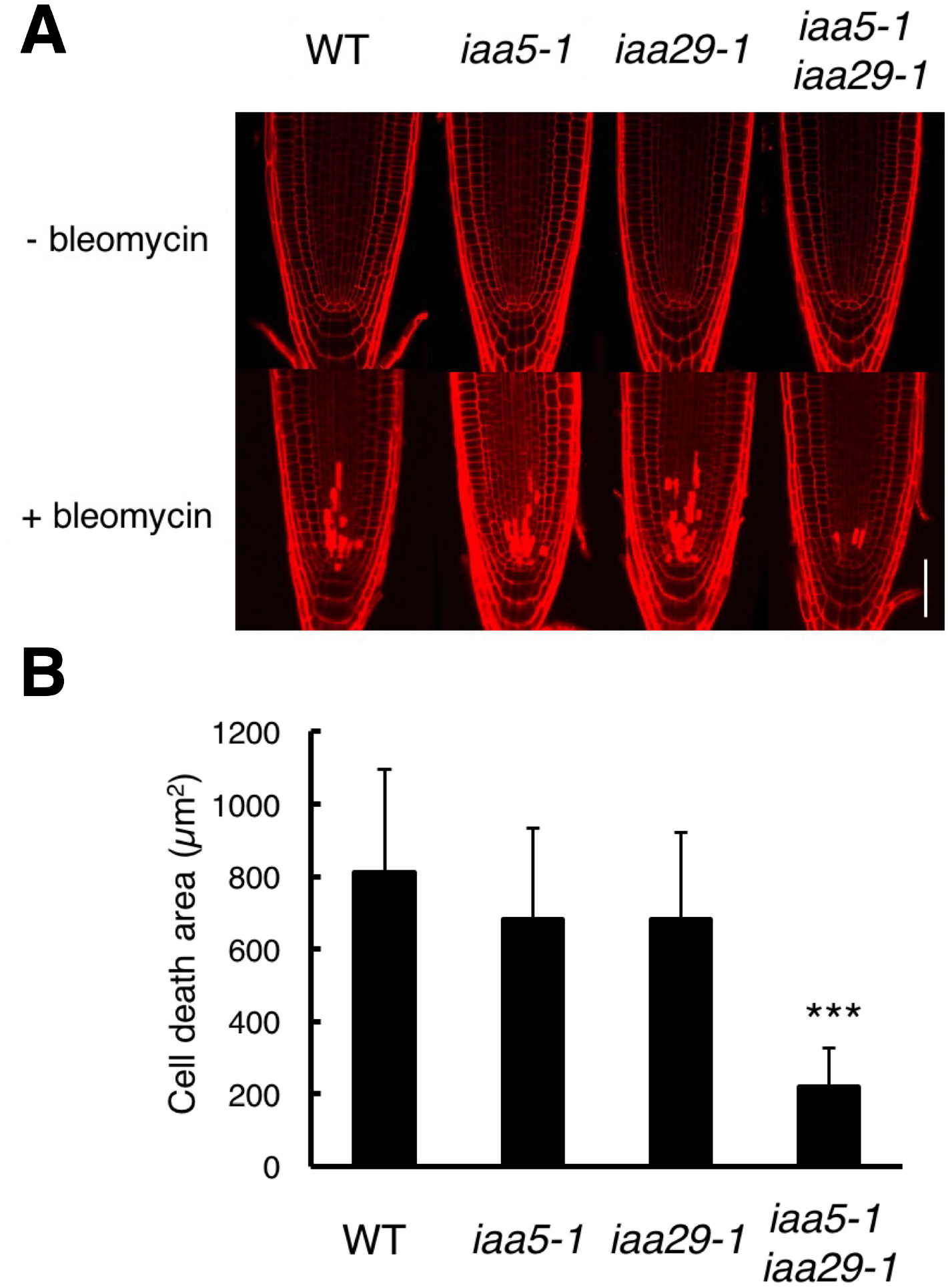
DSB-induced stem cell death is suppressed in the *iaa5 iaa29* double mutant. Five-day-old seedlings of WT, *iaa5-1, iaa29-1*, and *iaa5-1 iaa29-1* were transferred onto an MS medium supplemented with or without 0.6 μg/ml bleomycin, and grown for 24 h. (A) Root tips were observed after staining with PI. Bar = 100 μm. (B) The area of PI-stained dead cells in the root tip was measured using Fiji imaging software. Data are presented as mean ± SD (n > 20). Significant differences from WT were determined by Student’s *t*-test: ***, *P* < 0.001.

To further examine whether *IAA5* and *IAA29* locally induced in stem cells and their daughters indeed contribute to DNA damage-induced stem cell death, we expressed *IAA5*^*P59L*^, which encodes a stable form of IAA5 with a single amino acid substitution from proline to leucine in domain II (Ramos et al., 2001), in the *iaa5-1 iaa29-1* double mutant. *IAA5*^*P59L*^ was expressed under the *WOODENLEG* (*WOL*) promoter that is active in vascular stem cells and the stele (Petricka et al., 2012). We established two independent lines overexpressing *IAA5*^*P59L*^, which have no defect in the organization of vascular stem cells (Figure 5A and Supplemental Figure S5). Five-day-old seedlings were grown in a medium with or without bleomycin for 24 h, followed by PI staining. In the presence of bleomycin, the transgenic lines displayed stem cell death to the same extent as the wild-type (Figure 5, B and C), implying that a reduced level of cell death observed in *iaa5-1 iaa29-1* was recovered by *IAA5* expression in vascular stem cells and the stele. This result suggests that local *IAA5* induction promotes stem cell death in response to DSBs. Note that *pWOL:IAA5*^*P59L*^ *iaa5-1 iaa29-1* exhibited no stem cell death in the absence of bleomycin, indicating that *IAA5*^*P59L*^ overexpression alone cannot provoke cell death (Figure 5B).

**Figure 5.**
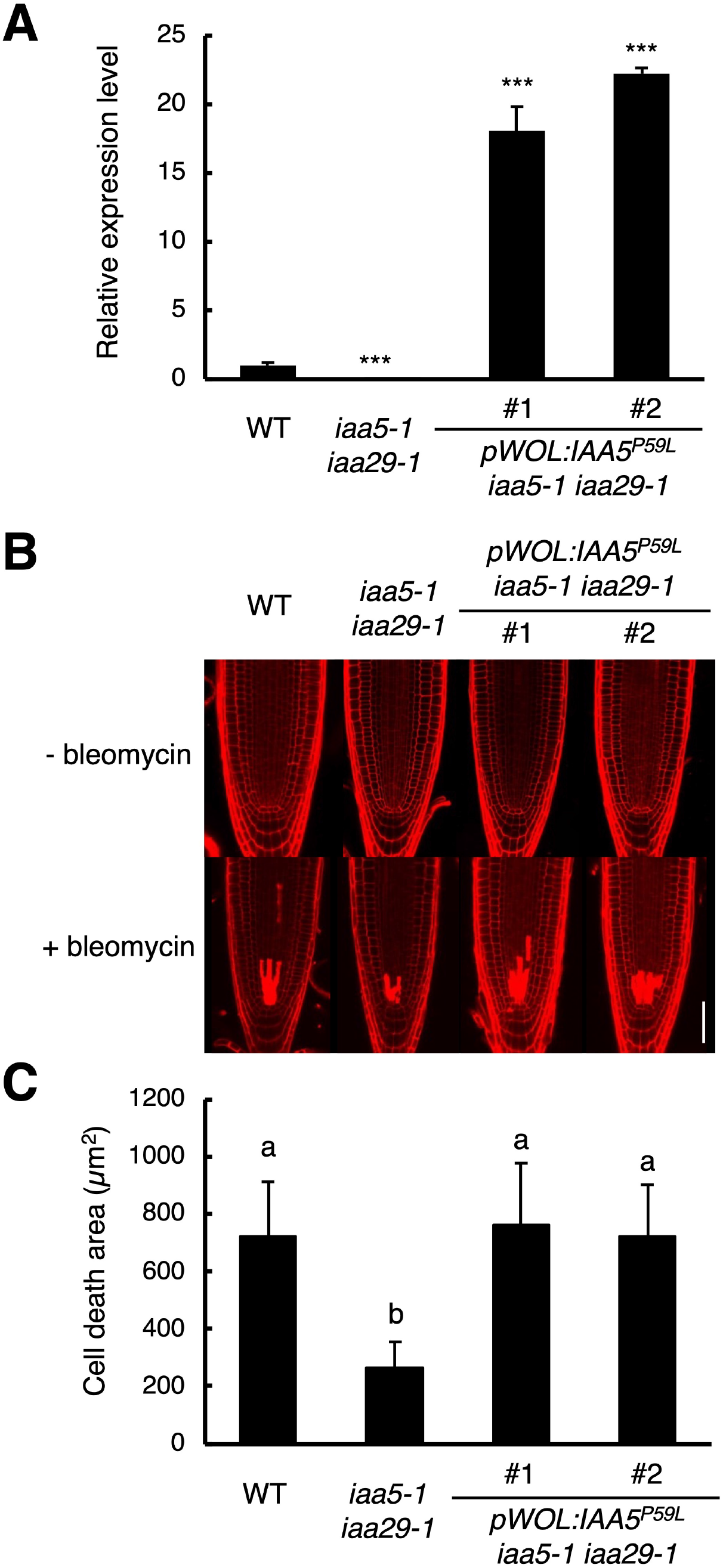
*IAA5*^*P59L*^ expression under the *WOL* promoter complements the *iaa5 iaa29* double mutant. (A) Transcript levels of *IAA5* and *IAA5*^*P59L*^. Total RNA was extracted from roots of five-day-old WT, *iaa5-1 iaa29-1*, and two independent lines of *pWOL:IAA5*^*P59L*^ *iaa5-1 iaa29-1* (#1 and #2). The mRNA levels of *IAA5* and *IAA5*^*P59L*^ were quantified by qRT-PCR using the *IAA5* primers and normalized to that of *ACTIN2*. They are indicated as relative values, with that of endogenous *IAA5* in WT set to 1. Data are presented as mean ± SD (n = 3). Significant differences from WT were determined by Student’s *t*-test: ***, *P* < 0.001. (B) Root tips of WT, *iaa5-1 iaa29-1*, and *pWOL:IAA5*^*P59L*^ *iaa5-1 iaa29-1*. Five-day-old seedlings were transferred onto an MS medium supplemented with or without 0.6 μg/ml bleomycin, and grown for 24 h. Root tips were observed after PI staining. Bar = 100 μm. (C) Cell death area in the root tip. The area of PI-stained cells was measured using Fiji imaging software. Data are presented as mean ± SD (n > 20). Bars with different letters are significantly different from each other (Student’s *t*-test: *P* < 0.05).

### IAA5 and IAA29 function in DSB-induced stem cell death independently from cytokinin-dependent reduction of auxin content

As described above, stem cell death was partially suppressed in bleomycin-treated *iaa5-1 iaa29-1* (Figure 4), suggesting that the IAA5- and IAA29-dependent reduction of auxin signaling is not the sole mechanism for cell death induction. We previously reported that DSBs upregulate cytokinin biosynthesis and inhibit the expression of several *PIN*s in the TZ, thereby weakening the downward auxin flow and decreasing the auxin level in the root tip (Takahashi et al., 2021). This suggests that the combinatorial mechanism reducing both the content and signaling of auxin plays a major role in cell death induction. To test this possibility, we used the *lonely guy 7* (*log7*) mutant defective in the final step of cytokinin biosynthesis (Kuroha et al., 2009). We previously demonstrated that in *log7*, the inhibition of *PIN* expression and the induction of stem cell death were suppressed in the root tip under DNA stress (Takahashi et al., 2021), suggesting that *log7* is an ideal material for genetic experiments. Our qRT-PCR data showed that bleomycin significantly induced *IAA5* and *IAA29* in *log7-1*, similar to wild-type (Figure 6A), supporting the idea that *IAA5* and *IAA29* induction is independent from enhanced cytokinin biosynthesis and is instead directly regulated by SOG1 (Figure 3). To investigate genetic interactions between *IAA5, IAA29*, and *LOG7*, we generated the triple mutant and measured cell death area after 24 h-bleomycin treatment. Stem cell death was suppressed in the *iaa5-1 iaa29-1* double mutant and *log7-1* to about 45% of that in the WT, whereas in the *iaa5-1 iaa29-1 log7-1* triple mutant, it reached less than 25% (Figure 6, B and C). This result suggests that DSBs induce *IAA5* and *IAA29* to reduce auxin signaling, as well as *LOG7* to decrease the auxin level, both of which are independently involved in stem cell death. When *IAA5*^*P59L*^ was overexpressed under the *WOL* promoter in the *iaa5-1 iaa29-1 log7-1* triple mutant, cell death area was recovered to the level of WT, but not to a lesser extent as observed in the *log7-1* mutant (Supplemental Figure S5 and S6). This indicates that the reduced cell death phenotype of *log7-1* as well as *iaa5-1 iaa29-1* was suppressed by *IAA5* overexpression; namely, *LOG7*-dependent *PIN* downregulation reduces auxin content, but eventually decreases auxin signaling in the root tip, thereby facilitating stem cell death.

**Figure 6.**
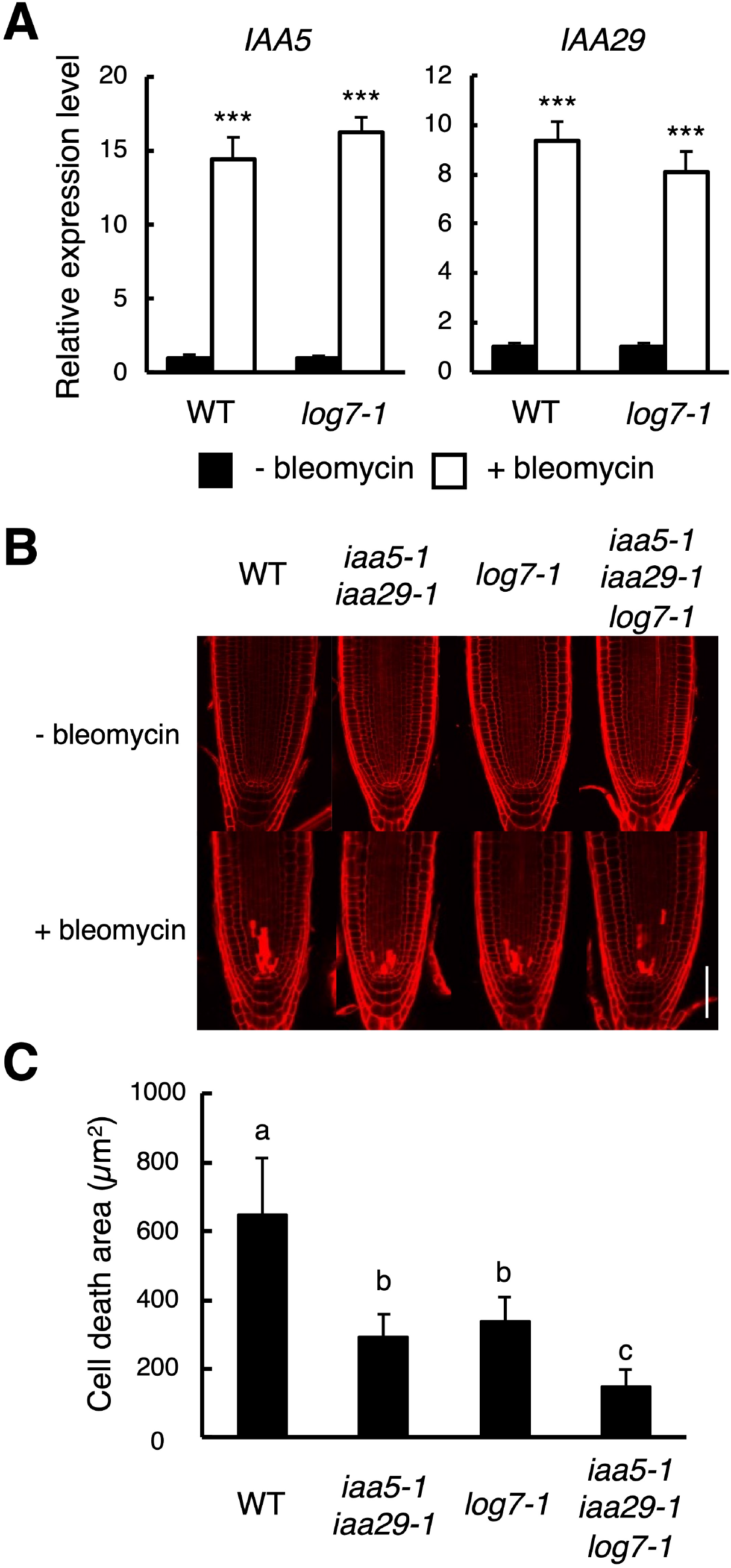
Additive effects of *iaa5 iaa29* and *log7* in suppressing DSB-induced stem cell death. (A) Transcript levels of *IAA5* and *IAA29* in *log7-1*. Five-day-old WT and *log7-1* seedlings were transferred onto an MS medium with or without 0.6 μg/ml bleomycin, and grown for 24 h. Total RNA was extracted from roots, and subjected to qRT-PCR. The mRNA levels of *IAA5* and *IAA29* were normalized to that of *ACTIN2*, and are indicated as relative values, with that of the control without bleomycin treatment set to 1. Data are presented as mean ± SD (n = 3). Significant differences from the control were determined by Student’s *t*-test: ***, *P* < 0.001. (B) Root tips of WT, *iaa5-1 iaa29-1, log7-1*, and *iaa5-1 iaa29-1 log7-1*. Five-day-old seedlings were transferred onto an MS medium supplemented with or without 0.6 μg/ml bleomycin, and grown for 24 h. Root tips were observed after PI staining. Bar = 100 μm. (C) Cell death area in the root tip. The area of the PI-stained cells was measured using Fiji imaging software. Data are presented as mean ± SD (n > 20). Bars with different letters are significantly different from each other (Student’s *t*-test: *P* < 0.05).

## Discussion

Plant tissues are generated post-embryonically, thus stem cells are essentially required for continuous organ formation as well as maintenance of organ homeostasis. Therefore, preserving genome integrity in stem cells is crucial for plant survival and reproduction. Plants, unlike animals in which severe DNA damage usually causes cell death, specifically induce cell death in stem cells, followed by stem cell replenishment through cell division at the organizing center, where DNA repair machineries are highly expressed (Heyman et al., 2013). This represents a sophisticated mechanism for maintaining genome integrity in plant stem cells. Indeed, stem cell death is an active response to DNA damage, which is mediated by the ATM/ATR–SOG1 pathway, in *Arabidopsis* (Fulcher & Sablowski, 2009; Furukawa et al., 2010). In this study, we revealed that SOG1 directly induces the two Aux/IAA family members *IAA5* and *IAA29* in vascular stem cells and their daughters, which are involved in DSB-induced stem cell death. We recently reported that SOG1 also upregulates *SHY2*, another Aux/IAA family member, indirectly through activating cytokinin biosynthesis (Takahashi et al., 2021). However, *SHY2* induction occurs at the boundary between the MZ and the TZ, represses a few *PIN* genes, and consequently reduces the auxin level over the MZ. This means that local induction of *IAA5* and *IAA29* further decreases auxin signaling specifically in stem cells and their daughters, thereby enabling stem cell replenishment. Considering that SHY2-mediated inhibition of downward auxin flow retards G2/M progression in the MZ (Takahashi et al., 2021), two distinct DNA damage responses, cell death in stem cells and G2 arrest in meristematic cells, are governed by Aux/IAA-mediated regulation of auxin signaling in the root tip.

A previous report demonstrated that, in tobacco-cultured cells, zeocin-induced DSBs were reduced by exogenous auxin treatment, while treatments with 2-(1H-Indol-3-yl)-4-oxo-4-phenyl-butyric acid (PEO-IAA), an inhibitor of the TIR1/AFB-mediated auxin signaling pathway, enhanced DSB accumulation (Hasegawa et al., 2018). This observation suggests that auxin has a role in protecting the genome from DNA damage. In addition, PEO-IAA treatments led to chromatin loosening, implying that auxin decreases chromatin accessibility and thus makes the genome more tolerant to DNA damage (Hasegawa et al., 2018). Based on this hypothesis, it is assumed that a dramatic reduction of auxin signaling causes chromatin relaxation in DNA-damaged stem cells, thereby providing hypersensitivity to DNA damage. However, gamma irradiation immediately induces DSBs without allowing time for reducing auxin signaling, nevertheless it triggers stem cell death as in the case of treatments with DSB-inducing agents (Furukawa et al., 2010). This suggests that downregulation of auxin signaling contributes to stem cell death after DSB induction, rather than sensitizing the genome to DNA damage.

Chen et al. (2017) reported that repressor-type Myb transcription factors (Rep-MYBs), which repress a set of G2/M-specific genes, are stabilized under DNA stress and lead to G2 arrest in *Arabidopsis*. In the *rep-myb* mutants, not only a reduction of cell division activity but also stem cell death is suppressed under DNA damage conditions (Chen et al., 2017), suggesting that G2 arrest is necessary to cause cell death. However, it is obvious that cell cycle arrest is not the only requirement for cell death induction; therefore, reduced auxin signaling is likely associated with cellular process(es) other than cell cycle regulation. Recent studies demonstrated that *Arabidopsis* Rep-MYBs form protein complexes called the DREAM/dREAM-like complex and coordinate cell cycle-dependent gene expression (Kobayashi et al., 2015; Lang et al., 2021). Interestingly, in *Drosophila* and vertebrates, the DREAM complex is known to regulate apoptosis as well as cell proliferation (Georlette et al., 2007; Sadasivam and DeCaprio, 2013). Therefore, a possible scenario is that, under DNA stress, auxin level affects the composition and/or activity of the DREAM complex, thereby fine-tuning cell cycle arrest and cell death; nonetheless, we cannot rule out the possibility that other factors are under the control of auxin signaling and control stem cell death. Further studies will reveal how DNA damage-induced cell death is confined to stem cells in plant meristems.

Lozano-Elena et al. (2018) demonstrated that DSB-induced stem cell death was suppressed in the *Arabidopsis* mutant of *BRASSINOSTEROID INSENSITIVE 1* (*BRI1*), which encodes one of the brassinosteroid receptors, indicating that brassinosteroid signaling promotes stem cell death. Another report showed that treatments with brassinolide, an active brassinosteroid, elevated the transcript levels of *IAA5* and some other *AUX/IAA* genes (Nakamura et al., 2006), raising a possibility that brassinosteroids enhance stem cell death by upregulating *Aux/IAA*s. On the other hand, brassinosteroids are also known to activate QC cell division to replenish stem cells under DNA stress (González-García et al., 2011; Vilarrasa-Blasi et al., 2014). Although the role of auxin in QC cell division remains elusive, the two hormones likely play a crucial role in coordinating stem cell death and QC cell division, thereby maintaining a proper number of stem cells under genotoxic stress. It is also probable that cell-to-cell communication may play an essential role in reorganizing the stem cell niche; for instance, stem cells send a death signal to the QC to activate cell division. Future studies will reveal how hormonal and intercellular signals are involved in stem cell renewal to preserve genome integrity in somatic plant cells.

## Materials and methods

### Plant materials and growth conditions

*Arabidopsis thaliana* (accession Col-0) was used in this study. Seeds were sown on MS plates [1/2 x MS salts, 1% sucrose, 0.5 g/L 2-(N-morpholino)ethanesulfonic acid (MES), and 1.2% phytoagar (pH 5.7)]. After incubation at 4°C for 2 days, plates were placed vertically under continuous light at 22°C. For DNA damage treatment, seedlings were transferred onto an MS medium containing 0.6 μg/ml bleomycin. *sog1-101* (Ogita et al., 2018), *iaa5-1* (Overvoorde et al., 2005), *iaa29-1* (Jiang et al., 2014), *log7-1* (Kuroha et al., 2009), and *pSOG1:SOG1-Myc* (Yoshiyama et al., 2013) were previously described. The *iaa5-1 iaa29-1* double and *iaa5-1 iaa29-1 log7-1* triple mutants were generated by crossing the single and double mutants. To generate the *promoter:GFP* fusion constructs, the 2-kb promoter fragments of *IAA5* and *IAA29* were PCR-amplified from *Arabidopsis* genomic DNA with the primer sets listed in Supplemental Table S1, and cloned into the pDONR P4-P1R entry vector (Invitrogen) by BP reaction according to the manufacturer’s instruction. Subsequently, LR reactions were conducted with the destination vector R4L1pGWB650 to make a fusion with *GFP* (Nakagawa et al., 2007). To generate *pWOL:IAA5*^*P59L*^, the *WOL* promoter and the *IAA5* gene were amplified by PCR with the primer sets listed in Supplemental Table S1, and cloned into the entry vectors pDONR P4-P1R and pDONR221 (Invitrogen), respectively, by BP reaction. To introduce the P59L mutation in *IAA5*, PCR was conducted using the primer set listed in Supplemental Table S1. An LR reaction was performed with the destination vector R4pGWB601 (Nakamura et al., 2010). All constructs were introduced into the *Agrobacterium tumefaciens* GV3101 strain, and stably transformed *Arabidopsis* plants were generated by the floral dip transformation method (Clough and Bent, 1998).

### qRT-PCR

Total RNA was extracted from roots with the Plant Total RNA Mini Kit (FAVORGEN). First-strand cDNA was synthesized from total RNA using ReverTra Ace^®^ qPCR RT Master Mix (Toyobo) according to the manufacturer’s instruction. qRT-PCR was performed with THUNDERBIRD SYBR qPCR Mix (Toyobo) using 100 nM primers and 0.1 μg of first-strand cDNAs. The primer sequences are listed in Supplemental Table S1. PCR reactions were conducted with the LightCycler 480 Real-Time PCR system (ROCHE) under the following conditions: 95°C for 5 min; 50 cycles at 95°C for 10 sec, at 60°C for 20 sec, and at 72°C for 30 sec.

### Microscopic observation

For PI staining, five-day-old roots were stained with 10 μM PI solution for 1 min at room temperature. Alternatively, to stain with SR2200, roots were soaked in an SR2200 solution [4% paraformaldehyde and 0.1% SCRI Renaissance 2200 (Renaissance Chemicals) in phosphate-buffered saline (PBS) (pH 7.4)] for 12 h at 4°C. Samples were washed with PBS and submerged in a ClearSee solution [10% xylitol, 15% sodium deoxycholate, and 25% urea] until roots became transparent. Root tips were observed under a confocal laser scanning microscope (Olympus, FluoView FV3000). Cell death area in the root tip was measured using Fiji software (https://fiji.sc).

### Chromatin immunoprecipitation (ChIP)-qPCR

ChIP was performed as described previously (Gendrel et al., 2005). Two-week-old seedlings grown in a liquid MS medium with gentle shaking (70 rpm) were treated with or without 0.6 μg/ml bleomycin for 12 h. Chromatin bound to SOG1-Myc proteins extracted from whole seedlings was precipitated with anti-Myc antibody (clone 4A6, Millipore). To quantify the precipitated chromatin, real-time qPCR was performed using the primer sets listed in Supplemental Table S1. PCR reactions were conducted with the LightCycler 480 Real-Time PCR system (ROCHE) under the following conditions: 95°C for 5 min; 60 cycles at 95°C for 10 sec, at 60°C for 20 sec, and at 72°C for 30 sec.

### Cell death induction by laser scanning

Five-day-old roots were laser-ablated with a confocal laser scanning microscope (Olympus, FV1000) equipped with the IR-LEGO unit (IR-LEGO 1000; Sigma-Koki). Cell death was induced by a 1480 nm laser generated by a high-power single mode CW Raman fiber laser (Model PYL-3-1480-M; IRE-Polus Group).

### Accession numbers

The genes used in this study have the following accession numbers: *IAA5* (AT1G15580), *IAA29* (AT4G32280), *SOG1* (AT1G25580), *WOL* (AT2G01830), *LOG7* (AT5G06300), and *Mut* (AT4G03870).

## Supplemental data

**Supplemental Table S1**. Primers used for cloning, qRT-PCR, and ChIP-qPCR.

**Supplemental Figure S1**. Expression levels of auxin-related genes in the presence of zeocin.

**Supplemental Figure S2**. Transcriptional responses of *IAA5* and *IAA29* to gamma irradiation and bleomycin.

**Supplemental Figure S3**. *IAA5* and *IAA29* expression in the root tip after 12 h-bleomycin treatment.

**Supplemental Figure S4**. No induction of *IAA5* and *IAA29* after laser irradiation.

**Supplemental Figure S5**. Cellular organization of vascular stem cells in the root tip.

**Supplemental Figure S6**. *IAA5*^*P59L*^ expression under the *WOL* promoter fully complements the *iaa5 iaa29 log7* triple mutant.

## Acknowledgments

We thank the *Arabidopsis* Biological Research Center (ABRC) for providing *Arabidopsis* mutants.

